# Time-dependent mnemonic vulnerability induced by new-learning

**DOI:** 10.1101/281261

**Authors:** Fengtao Shen, Yixuan Ku, Jue Wu, Yue Cui, Jianqi Li, Zhaoxin Wang, Huimin Wang, Sze Chai Kwok

## Abstract

Reactivation renders consolidated memory labile again, and the ensuing temporary reconsolidation process is highly susceptible to mnemonic modification. Here, we show that memories in such an unstable state could be reprogrammed by sheer behavioral means, bypassing the need for pharmacological intervention. In two experiments using a “face-location associationc” paradigm in which participants experienced a “Learning – New-learning – Final-test” programme, we demonstrate that reactivated memory traces were robustly hampered when the new learning was strategically administered within a critical 20-minute time window. Using fMRI, we further advance our theoretical understanding that this lability can be mechanistically explained by the differential activation in the hippocampal-amygdala memory system implicated by the new-learning whereas the mnemonic intrusion caused by newly learned memories is efficaciously reconciled by the left inferior frontal gyrus. Our findings provide important implications for educational and clinical practices in devising effective strategies for memory integration.

## 1 Introduction

Memory recall is constructive in nature and the mere act of recalling a memory renders it labile and highly susceptible to modification (Nader, Schafe et al. 2000, Lee, Everitt et al. 2004, Lee, Di Ciano et al. 2005, Alberini and LeDoux 2013, Lee, Nader et al. 2017, Scully, Napper et al. 2017). While emotional factors are known to exert effects on reconsolidation of emotional declarative memories (Schwabe and Wolf 2009, Strange, Kroes et al. 2010, Chan and LaPaglia 2013), controversies still surround reconsolidation theories on non-emotional declarative memories. Empirically, post-retrieval manipulations gave rise to inconclusive patterns of results, with some studies showing such manipulation can induce update (Hupbach, Gomez et al. 2007, Hupbach, Gomez et al. 2009, Forcato, Rodriguez et al. 2010), forgetting (Forcato, Burgos et al. 2007), extinction (Nader, Schafe et al. 2000, Schiller, Monfils et al. 2010, Agren, Engman et al. 2012), or enhancement (Coccoz, Maldonado et al. 2011, Coccoz, Sandoval et al. 2013), whereas another set of studies revealing no observable effect (Cammarota, Bevilaqua et al. 2004, Debiec, Doyère et al. 2006, Hupbach, Hardt et al. 2008, Forcato, Argibay et al. 2009, Hupbach, Gomez et al. 2011, Gershman, Schapiro et al. 2013). These previous studies indicate that manipulations after reactivation would induce multiple, and at times conflicting, effects under different conditions (Nader, Schafe et al. 2000, Pedreira, Perez-Cuesta et al. 2002, Walker, Brakefield et al. 2003, Debiec, Doyère et al. 2006, Forcato, Argibay et al. 2009, Sederberg, Gershman et al. 2011, Sevenster, Beckers et al. 2012), it was thus important to characterize these contributory factors. Specifically, reconsolidation is known to be time-sensitive. The presence of this time-dependence in humans has been coarsely derived from studies utilizing either one of the two extreme reactivation-intervention intervals: either too short such that the reconsolidation was still ongoing (e.g., 5 or 10 minutes, (Forcato, Burgos et al. 2007, Forcato, Argibay et al. 2009, Schiller, Monfils et al. 2010, Agren, Engman et al. 2012)), or too long such that the reconsolidation had concluded before the intervention began (e.g., 6 or 10 hours, (Forcato, Burgos et al. 2007, Schiller, Monfils et al. 2010, Agren, Engman et al. 2012)). Here we investigated the detailed temporal characteristics of reconsolidation of declarative memory using gradient-like post-reactivation delays.

In light of the controversies surrounding theories on the reconstructive nature of declarative memories, we evinced that human associative memories can be exquisitely rendered labile by newly-acquired memories within a critical time-window. Using a face-location association learning paradigm, human participants were made to experience acquisition, test of associative-learning, reactivation, new-learning, and final-test across three consecutive days. In a behavioral experiment (Fig. 1A, upper panel), participants encoded 30 face-location associations on Day 1 (day1-Acquisition) and following a 24-hour retention period, they were then divided into 5 groups and asked to recall the associations they had acquired previously on day1 (day2-Reactivation). Importantly, the four different groups of participants received a critical time-dependent new-learning manipulation (i.e., acquiring a new location associated with the original 30 faces) whilst a fifth group acted as a control group and did not receive any new-learning. The day2-New-learning served a critical interventional purpose, aiming at interfering the originally acquired memories during reconsolidation. On the third day (day3-Final-test), these participants were required to recall again the face-location associations they had learned on day1-Acquisition. We revealed the new-learning that occurred right after reactivation significantly diminished the memory of the originally learned associations in a time-dependent manner.

**Figure 1.**
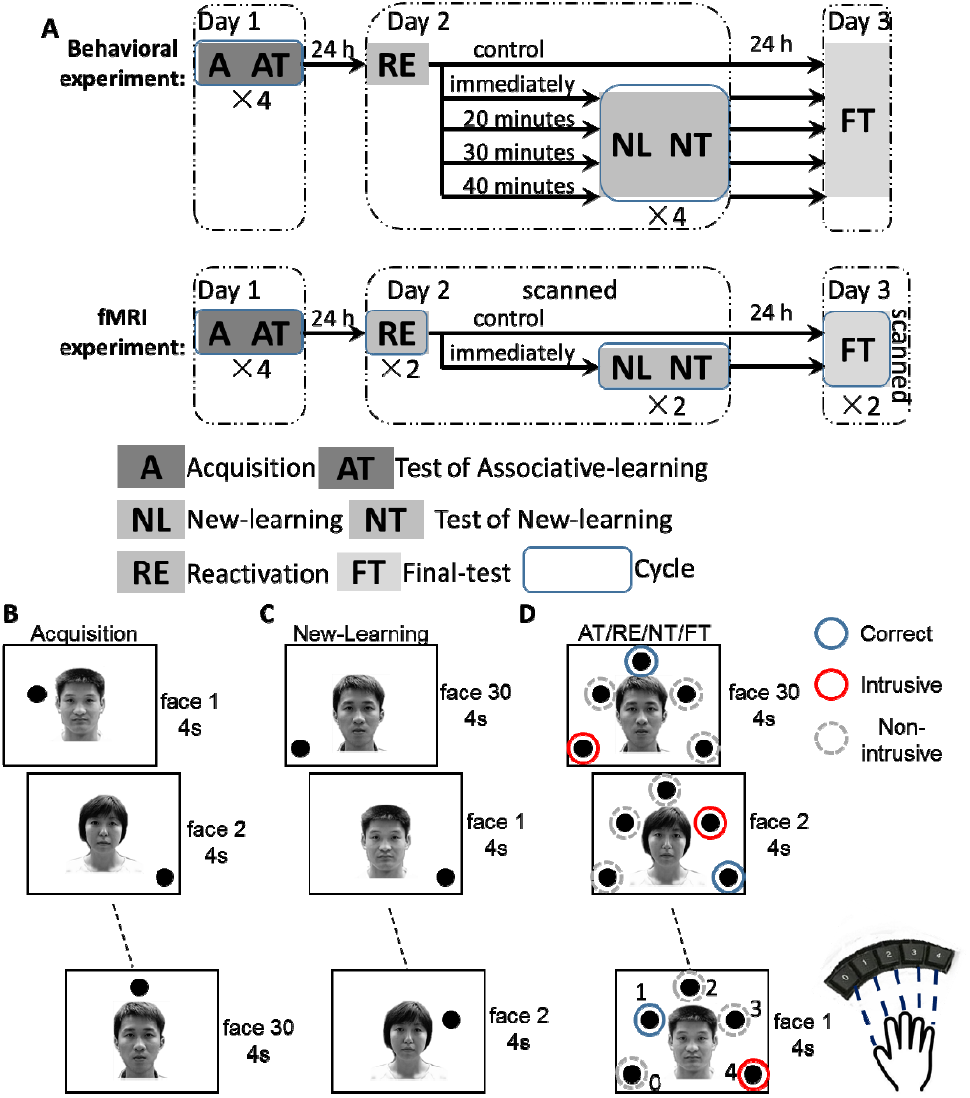
Paradigm Overview. (A) Experimental overview for behavioral and fMRI experiments. There were five and two groups in the two experiments, respectively. Each of the experiments, spanning across 3 daily sessions, consisted of four stages: Acquisition, Test of Associative-learning (Day 1), Reactivation, New-learning, Test of New-learning (Day 2), and Final-test (Day 3). On Day 1, the subjects acquired a set of face-location associations (Acquisition). On Day 2, they were first asked to recall original associations (Reactivation) and were then divided into 4 experimental groups and one control group. After variable delays (i.e., 0’, 20’, 30’, and 40’), they learned another set of associations of linking a new location to each of the original faces (New-learning). The control group did not receive any new learning. Finally, on Day 3, these subjects were asked to recall the originally learned associations which they had acquired on Day 1 (Final-test). The participants in the fMRI experiment were scanned on Days 2 and 3. The cycle “×4” and “×2” denote the numbers of repetition in each of the tests. (B) Original learning (Acquisition) consisted of 30 face-to-location associations. On each trial, a unique face was presented together with a location (out of five possible locations) on the screen for 4 s. The participants were instructed to memorize the associations. Their memories were then tested with Tests of Associative-learning. No feedback was given. (C) Importantly, using the identical procedure, on Day 2, 30 new associations were acquired *de novo* by the participants in the New-learning stage. (D) In Test of Associative-learning (AT), Reactivation (RE), Test of New-learning (NT), and Final-test (FT) stages, on each trial, the participants were required to indicate the correct location matched to each of the faces by pressing a 5-button keypad. In the Final-test, each response was classified into either a Correct response (blue discs), an Intrusive error (red discs), or a Non-intrusive error (grey discs). The colored discs, the face ID numbers and the location numbers (0-4) were not shown in the actual experiment. The order of face-presentation was randomized within and across participants in all stages.

**Table 1.**
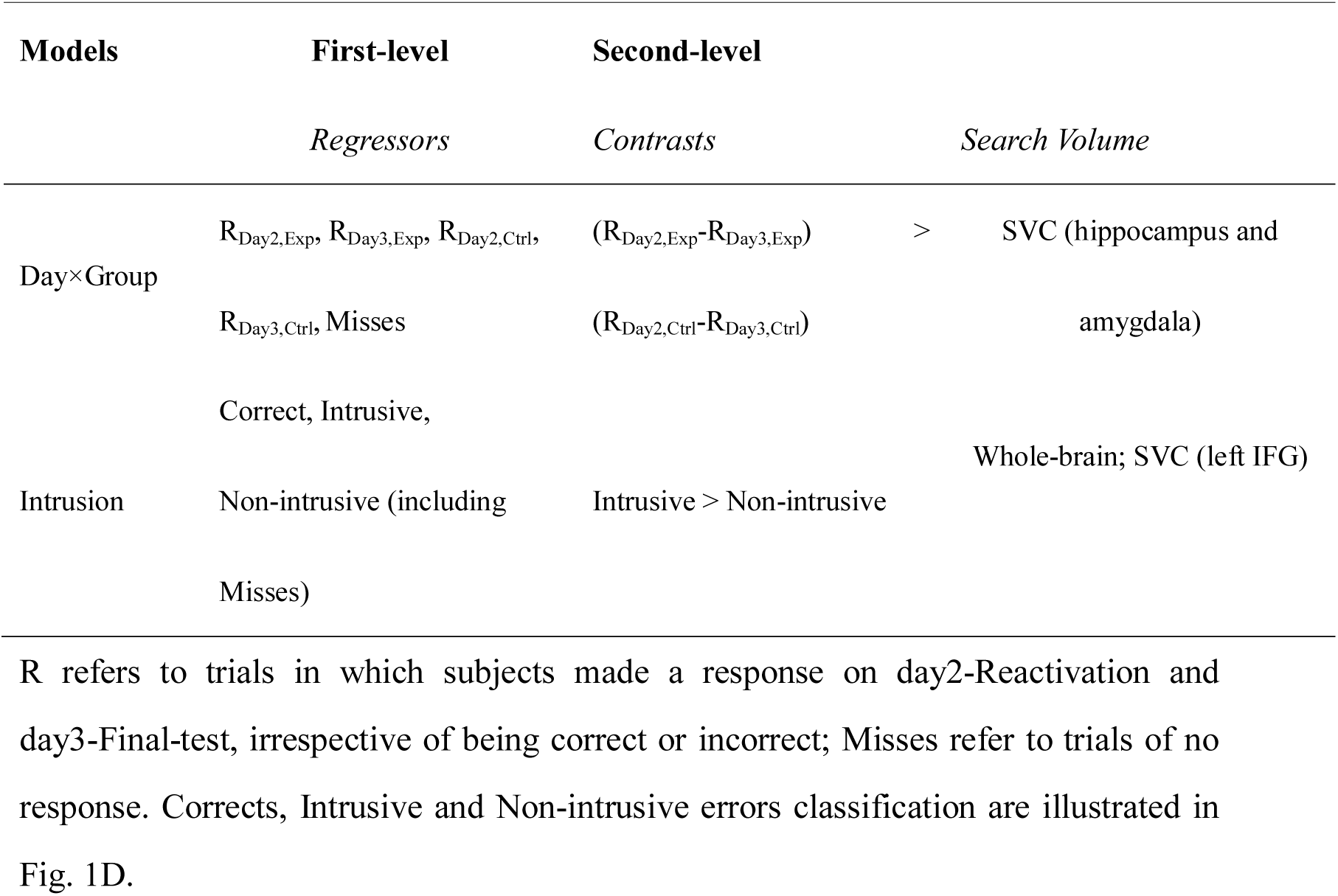
Summary of all 1st-/2nd-level analyses and contrasts for both models.

To elucidate the behavioral effects induced by the new-learning and the neural underpinnings of the reconsolidation processes, we replicated the behavioral experiment with a new group of participants performing a corresponding experiment while their blood-oxygen-level-dependent (BOLD) activity was measured. We probed, at a macro-anatomical level, in which regions might lie the influence of the new-learning on the reconsolidation of non-emotional episodic memory (i.e., how new-learning affected the originally learned memory) and how the intrusive effects thereby induced by the newly-learned associations might manifest neurally. In this fMRI experiment we included only one experimental group, which began their new-learning immediately after reactivation on Day 2 (Fig. 1B-D). We employed fMRI to unravel the mechanisms underlying the processes of integrating new information into consolidated memories during reconsolidation.

## 2 Method

The entire study consisted of one behavioral and one fMRI experiment. Each consisted of four experimental sessions across three days. The separation between days was strictly controlled to be 24 hours (Fig. 1).

### 2.1 Subjects

151 participants took part in the behavioral experiment proper (four experimental groups n = 28 each; control group n = 39) and 46 participants took part in the fMRI experiment (experimental group n = 18; control group n = 28). All of them were recruited from the East China Normal University (17 – 30 years old, mean = 22.05±2.51, SD, 26 males). All had normal or corrected-to-normal vision and reported regular nocturnal sleep and no history of any neurological, psychiatric or endocrine disorder. The participants received monetary compensation for their participation. Written informed consents were obtained from all participants and the study was approved by the University committee on Human Research Protection (UCHRP) at East China Normal University. An additional 60 participants (37 and 23 for the behavioral and fMRI experiments, respectively) were recruited but were not invited to enter the subsequent sessions because their performance accuracy was below 40% in the last round of the Test of Associative-learning on Day 1.

### 2.2 Stimuli

60 grayscale front-facing faces of neutral expression from unfamous volunteers (30 males) were selected from CAS-PEAL-R1 database (http://www.jdl.ac.cn/peal/index.html). These were divided into two sets of 30 faces each (each set consisted of 15 males and 15 females). One set was used in the behavioral and fMRI experiments, in which both day1-Acquisition and day2-New-learning employing the same set of 30 faces (Fig. 1). The second set was specifically used in the day2-New-learning phase for a control experiment wherein the 30 faces used in the new-learning were different from those used in the original learning session (Supplementary Fig. S1).

### 2.3 Behavioral experiments and analysis

The behavioral experiment investigated how the time interval between memory recall (i.e., reactivation) and interference (i.e., new-learning) affects memory reconsolidation. Four gradient-like time intervals between memory recall and interference were chosen: 0, 20 minutes, 30 minutes, and 40 minutes (four experimental groups). To obtain a reliable baseline for comparison, a control group was included in which no interference was applied (control group).

In a face-location association learning paradigm, the participants first familiarized themselves with the 30 faces on Day 1 (familiarization session) by viewing these faces passively. Each face was presented at the center of the screen for 3 s and separated by a jittered inter-trial interval of 2-4 s (mean = 3s). The whole set of 30 faces was presented three times in a randomized order.

Following the familiarization phase, the participants were then asked to memorize 30 face-location associations (day1-Acquisition; A), involving each face being paired with one of five location points on the screen. They were allowed 4 s to learn each pairing (Fig. 1B). Immediately after each acquisition of the 30 face-location pairings, a memory test ensued (Test of Associative-learning, AT, Fig. 1). On each test trial, the face cue and all five location points were presented together, and the participants were asked to indicate within 4 s which location disc was originally paired with the face in the Acquisition stage by pressing the button corresponding to the target location using an MRI-compatible keypad (see cartoon in Fig. 1D). This Acquisition – Test of Associative-learning procedure was repeated four times with the set of face-location associations presented in a new randomized order in each cycle. The trials were separated by jittered inter-trial intervals of 3-7 s (mean = 5s) and no feedback was given.

On Day 2, the participants were asked to recall their memory of the previously learned face-location associations by identifying the target location that was associated with a given face (day2-Reactivation; RE, Fig. 1). A New-learning procedure was then administered aiming to interfere the processes of memory reconsolidation. The participants were asked to learn to associate the originally-learned faces with a new target location (i.e., learning new face-location associations, Fig. 1C). This New-learning session consisted of four cycles of New-learning (NL) and Test of New-learning (NT).

In order to pinpoint the temporal characteristics of interference on memory reconsolidation, four temporal intervals, namely 0’, 20’, 30’, and 40’, between the day2-Reactivation and New-learning were administered separately to the four experimental groups. During these post-reactivation intervals, the participants listened to light music without having to perform any task.

On Day 3, the participants recalled the face-location associations they had acquired on Day 1 (day3-Final-test; FT), identifying the target locations that were associated with given faces from Day 1.

A mixed 5 (between-group factor, four experimental conditions and control condition) × 3 (within-group factor: Day1, Day2 and Day3) analysis of variance (ANOVA) was applied on percentage correct data from the behavioral experiment. Analogously, a mixed 2 (Group Exp. and Ctrl.) ×3 (Day1, Day2 and Day3) ANOVA was applied on the data from the fMRI experiment.

Moreover, to account for inter-subject variability, the within-subjects correct rates were normalized to obtain relative correct rates using the following equations,

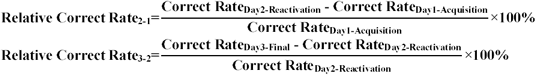

The within-subjects relative Correct Rate_2-1_ reflects the memory decay after Day1-Acquisition before Day2-Reactivation, whereas the relative Correct Rate_3-2_ reflects the memory change due to the New-learning intervention.

### 2.4 Classification of correct, intrusive and non-intrusive responses

During the Final-test session, the participants were instructed to respond to the target location as they learned in the acquisition on Day 1. Since the experimental groups experienced new-learning on Day 2, there were three categories of responses in the day3-Final-test. If the response was correctly matched with acquisition, it was a correct hit. If it was incorrectly matched with the location they acquired in the new learning on Day 2, it was classified as an intrusive error. Responses made to the other three locations would be non-intrusive errors (Fig. 1D). We compared the difference between the correct and the intrusive proportions among the groups. If the correct rate/intrusive ratio was not significantly different between the experimental groups and the control group, then we would infer that new learning did not cause any significant effect. By contrast, if there were significant differences in the correct rate/intrusion ratio between the groups, we would conclude that the new learning might have disrupted the original-memory more severely in the experimental group(s).

### 2.5 Control experiment: Effectiveness of content-similarity in memory intervention

In declarative memories, content similarity shared between the acquisition and new-learning material is a key factor for effective intervention as only similar new materials were found to induce memory update, disruption or enhancement via reconsolidation (Forcato, Burgos et al. 2007, Hupbach, Gomez et al. 2007, Coccoz, Maldonado et al. 2011, Forcato, Rodriguez et al. 2011). We hypothesized that material used in the post-reactivation intervention has to be similar enough to those used in the acquisition to cause any discernible effect on the reconsolidation processes. To test this prediction, we ran an additional control experiment in which we utilized new and unencountered faces as the post-reactivation new-learning material (i.e., new faces to be paired up with the original locations).

### 2.6 MRI acquisition and preprocessing

Participants were scanned in a 3T MRI scanner (Trio Tim, Siemens) with a quadrature volume head coil at the Shanghai Key Laboratory of Magnetic Resonance. Thirty-three slices of functional MR images were acquired using a gradient EPI sequence (EPI volumes per run = 192, FOV = 210×210 mm^2^, matrix = 64×64, in-plane resolution = 3.75×3.75 mm^2^, thickness = 4 mm, without gap, repetition time = 2 s, echo time = 30 ms, flip angle = 90°), covering the entire brain. A high-resolution structural image for each participant wasp also acquired using 3D MRI sequences for anatomical co-registration and normalization (FOV = 256×256 mm^2^, matrix = 256×256, slice thickness=1 mm, without gap, repetition time = 2530 ms, echo time = 2.34 ms, flip angle = 7°).

SPM8 (Wellcome Department of Cognitive Neurology, London, UK; http://www.fil.ion.ucl.ac.uk/spm/) was used for data processing. For each participant, the functional images were realigned to correct for head movements. The structural image was co-registered with the mean EPI image, then segmented and generated normalized parameters to MNI space. Functional images were then normalized to the MNI space using these parameters, re-sampled to 2 mm isotropic voxel size and then spatially smoothed using an isotropic Gaussian kernel of 8 mm FWHM (full-width half-maximum). High-pass temporal filtering with a cut-off of 128 s was performed to remove low-frequency drifts.

### 2.7 fMRI data analysis

The fMRI experiment examined the neural correlates underlying the several aspects elicited by the new-learning interference on memory reconsolidation. The same face-location association learning paradigm as in the behavioral experiment was adopted. We implemented two Reactivation sessions on Day 2 and also two Final-test sessions on Day 3 to ensure a decent volume of data to be collected for the fMRI analyses. Across the two sessions, we collected data for 60 test trials (i.e., two repetitions of the complete set of the 30 face-location pairs). Informed by the behavioral experiment that the memory reconsolidation processes were susceptible after a 0’-delay, we accordingly targeted at the 0’ condition here. We included one control group, which received no new-learning after reactivation, for comparison. Trials were separated by jittered inter-trial intervals of 3-7 s (mean = 5s) and 18 4-s blank trials were included as baseline measurement. Each of the New-learning runs lasted for 10 min and each of the Reactivation/Final-test runs lasted for 6 min (Fig. 1C-D). The fMRI data for the day2-New-learning were not included for analysis.

Two sets of analyses (Day×Group model and Intrusion model) were carried out using a general linear model (**Error! Reference source not found.**). Statistical inference was based on a random effects approach, which comprised first-level analyses estimating contrasts of interest for each subject and second-level analyses for statistical inference at the group level with non-sphericity correction. For both models, in the first-level, each of the 60 test trials was modelled with a canonical hemodynamic response function time-locked to the trial onset as an event-related response with that trial’s duration (mean duration = 2466 ms). The design matrix included six head motion regressors to remove the residual effects of head motion. The blank trials were not modelled. The estimated parameters values were used for the second-level group analysis.

The first model (Day×Group model) sought to identify brain areas that activated during reactivation and final-test. This allowed us to calculate the interaction effect between the two factors for finding any evidence of episodic memory reconsolidation. In the first-level analysis, the model included five regressors: R_(Day2,Exp)_, R_(Day3,Exp)_, R_(Day2,Ctrl)_, R_(Day3,Ctrl)_, Misses, reflecting the responses of the experimental and control groups in day2-Reactivation and day3-Final-test. For the group-level analysis, the single-subjects contrast images for the 2 experimental conditions (i.e., “Day2/Day3” trials, averaged across the two fMRI-runs) for each of the two groups were entered into a mixed design ANOVA with “Day” as the within-subject variable and “Group” as the between-groups variable. The random effects analysis consisted of an ANOVA assessing the significance of Delta *T*-covariate at the group level. The statistical threshold was set to *P*-FWE=0.05, whole brain corrected at peak level (cluster size estimated at *P*-unc. = 0.005). With our *a prior* prediction, we performed small volume correction (SVC) using a functional mask derived from subsequent memory effects as the volume of interest (covering the hippocampus and the amygdala, (Kim 2011)).

The second model (Intrusion model) concerned responses during the final-test, specifically investigating how new learning affected the original memory trace during the reconsolidation process. The first-level model included three regressors obtained from the day3-Final-test, reflecting the three types of the responses (correct, intrusive or non-intrusive). Six motion regressors were also included. For the group-level analysis, the single-subjects contrast images for the 3 experimental conditions (i.e., “correct/intrusive/non-intrusive” trials, averaged across the two fMRI-runs) were entered into an ANOVA. The statistical threshold was set to *P*-FWE=0.05, whole brain corrected at peak level (cluster size estimated at *P*-unc. = 0.005). The random effects analysis consisted of a one-sample t-test assessing the significance of Delta *T*-covariate at the group level. Specifically, the “Intrusive > Non-intrusive” contrast revealed a cluster in the left inferior frontal gyrus. We accordingly extracted the beta estimates of the left IFG from each subject using Marsbar and correlated these beta estimates with the proportion of correct responses and the proportion of intrusive errors separately.

## 3 Results

### 3.1 Behavioral results: Main experiment

We revealed compelling evidence in support of the existence of reconsolidation. In the behavioral experiment, the Day × Group repeated measures ANOVA on percentage correct showed a strong “Day × Group” interaction effect (F _(8, 292)_ = 7.26, *P* < 0.001, Fig. 2A, Supplementary Table S1). We then ran two separate ANOVAs and found the group differences were only in the day3-Final-test, (F _(4, 146)_ = 5.11, *P* = 0.002, Fig. 2A, Supplementary Table S1) but not in day2-Reactivation (F _(4,146)_ =0.39, *P* = 0.81). In order to account for individual variances, we normalized the percentage correct data and re-ran ANOVAs on these synthetic, more sensitive indices. A 2 (correct rate_2-1_; correct rate_3-2_) x 5 (Group) repeated measures ANOVA equally showed a strong interaction between the factors (F _(4,146)_ = 6.00, *P* < 0.001). Two separate ANOVAs showed that the interaction was driven by a main effect in relative correct rate_3-2_ between Days 2 and 3, confirming that the significant between-group differences were specifically caused by new-learning (relative correct rate_3-2_: F _(4, 146)_ = 9.75, *P* < 0.001, Fig. 2B right, Supplementary Table S2) but not before new-learning (relative correct rate_2-1_: F _(4, 146)_ = 0.39, *P* = 0.81, Fig. 2B left).

**Figure 2.**
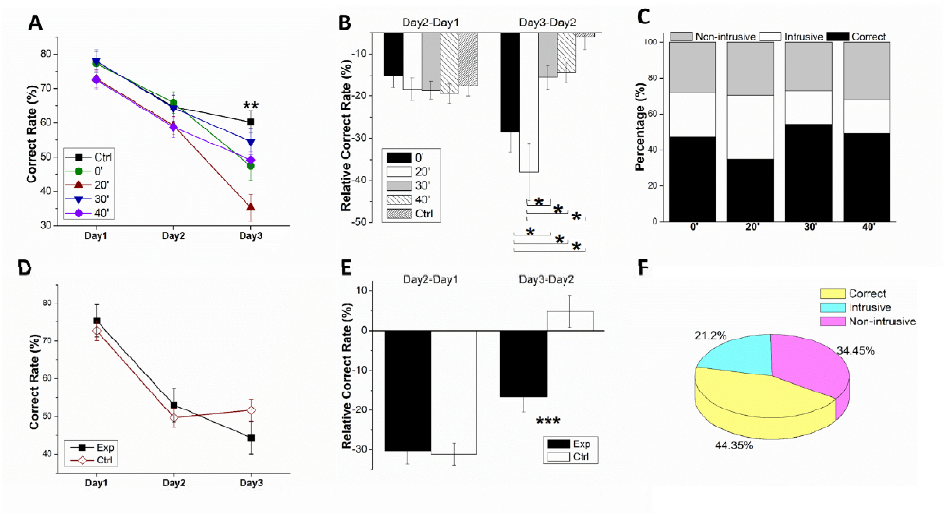
Behavioral results of both experiments. (A) Memory performance plotted as a function of days in the behavioral experiment. Memory of the face-to-location associations diminished in all five groups gradually across days but a main effect of Group was found on Day 3 at the Final-test. (B) The reduction in memory in the behavioral experiment was plotted for relative correct rate_2-1_ and relative correct rate_3-2_, respectively. There was no difference in Group for relative correct rate_2-1_ but there was a significant interaction for the relative correct rate_3-2_. Post-hoc tests confirmed that the memory for the Group 0’ and 20’ decreased far more drastically than Group 30’, 40’ and the control group. (C) The intrusive proportion of Group 20’ was significantly larger than other groups. (D) Behavioral result in the fMRI experiment was consistent with that of the behavioral experiment. Both Group Exp. and Ctrl. performed similarly on Day 2. But the performance of the Group Exp., who had received post-reactivation New-learning on Day 2, diminished far more severely than Group Ctrl. at the Final-test. (E) Using a relative measure, in the fMRI experiment, there was no Group difference in the relative correct rate_2-1_, but Group Exp. was significantly more impaired than Group Ctrl. in the relative correct rate_3-2_. (F) The intrusive proportion of Group Exp. in the fMRI experiment (21.2%) was similar as Group 0’ in the behavioral experiment (cf. leftmost bar in Fig. 2C). Error bar denotes standard error of the means. * *P*<0.05, ** *P*<0.01, *** *P*<0.001.

Motivated by previous findings on the time-dependence of post-retrieval manipulations (Forcato, Burgos et al. 2007, Schiller, Monfils et al. 2010, Agren, Engman et al. 2012, Chan and LaPaglia 2013), we then tested the hypothesis that there would be a critical time-window for the observable post-reactivation reconsolidation. As expected, the difference in the relative correct rates between Day 2 and Day 3 for Group 0’ and 20’ were significantly lower that other three groups (all *P*s < 0.05, LSD multi-comparison, Fig. 2B), indicating the influence of new-learning was indeed highly time-dependent.

It has been reported that new-learning could produce an intrusive effect to our memories by replacing the original memories in a specific retrograde manner (Hupbach, Gomez et al. 2007, Hupbach, Gomez et al. 2009, Schiller, Monfils et al. 2010). In view of this, we tested for the intrusive effect in the current context. On each trial, there were five location points; each of which could be a potential choice. Operationally, for the experimental groups, at day3-Final-test, responses made to the target location would be a hit, responses made to the newly-learned location would be an intrusive error, whereas responses made to any of the other three locations would be a non-intrusive error (see Methods). The intrusive proportion of Group 20’ was significantly higher than other three groups (all *P*s < 0.05, Fig. 2C, Supplementary Fig. S2, Supplementary Table S3), whereas these intrusive errors in the other three groups did not differ. Interestingly, in Group 20’, the intrusive proportion did not differ from the correct rate, while in other three groups the correct rates were significantly higher than the intrusive proportions (Supplementary Fig. S2, Supplementary Table S3), implying the new-learning might have caused differential effects on Group 0’ and 20’.

We further analyzed these intrusive effects in all experimental groups. Interestingly, the quantity of intrusive errors in the 20’ condition is significantly higher than those in the other conditions (F _(3,108)_ = 5.08, *P* = 0.003; LSD multi-comparison: *P* _(20’>0’)_ = 0.035*, *P* _(20’>30’)_ = 0.001**, *P* _(20’>40’)_ = 0.001** vs. *P* (0’>30’) = 0.21, *P* _(0’>40’)_ = 0.23, *P* _(30’>40’)_ = 0.96), indicating the intrusive effects induced by new-learning following different post-reactivation delays are differential.

### 3.2 Control experiment results: Effective manipulation requires high content-similarity between acquisition and intervention

In this control experiment, the new-face-learning caused no effect on reconsolidation. We ran a 3 × 2 repeated measures ANOVA (Day×Group) on percentage correct and found neither a group main effect nor an interaction effect (group main effect: *F* _(1,2)_ = 1.49, *p* = 0.24; interaction: *F* _(2,2)_ = 1.29, *p* = 0.29; Supplementary Fig. S1). We conducted the post-hoc tests regardless and confirmed there were no group differences in day2-Reactivation (*t* _(1, 20)_ = 0.47, *p* = 0.65) or day3-Final-test (*t* _(1, 20)_ = −0.22, *p* = 0.82), nor in the relative correct rate_3-2_ between Days 2 and 3 (*t* _(1,) (20)_ = −1.36, *p* = 0.19). These indicate that new-learning using “newfaces” was ineffective in causing interference in the memory traces during reconsolidation.

### 3.3 fMRI experiment results

We have thus far established in the behavioral experiment that new-learning following reactivation did intrude into the already encoded, yet labile memories, and produce overt changes in terms of memory behavior. We then tap into the rather complicated and unresolved mechanisms of reconsolidation by means of functional imaging. We replicated these behavioral patterns in the fMRI experiment with a new group of participants. A 2×2 repeated measures ANOVA (Day×Group) showed an interaction effect (F _(2, 88)_ = 6.86, *P* = 0.002, Fig. 2D). The performance for the experimental group was significantly lower than that of control group in the relative correct rate_3-2_ (t _(44)_ = −3.65, *P* < 0.001, Fig. 2E right) but not in the relative correct rates_2-1_ (t _(44)_ = 0.18, *P* = 0.860, Fig. 2E left).

To look into the neural correlates, we ran a “Day×Group” model to test for the interaction between Day and Group to look for the effects of new-learning on original memory. Specifically, the interaction term (R_Day2,Exp_-R_Day3,Exp_) vs. (R_Day2,Ctrl_-R_Day3,Ctrl_) revealed activation of left hippocampus and right amygdala (Fig. 3). Both regions yielded significant activation (hippocampus: peak *P*-svc = 0.049; amygdala: peak *P*-svc = 0.037) with small volume correction (SVC) (volume-of-interest obtained from a subsequent memory effects contrast: remembered vs. forgotten)(Kim 2011). Notably, the amygdala has been known to be related to emotional processes especially by those that are involved in fear and threat memory reconsolidation (Agren, Engman et al. 2012, Schiller, Kanen et al. 2013). However, in the present setting, considering our paradigm did not contain any emotional factors, the right amygdala was implicated regardless.

**Figure 3.**
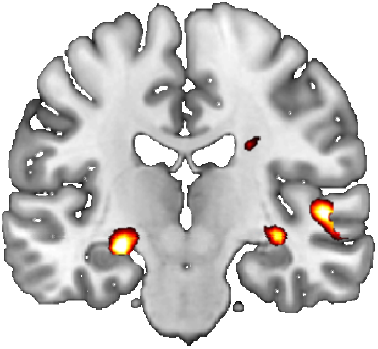
Neural correlates associated with the impact of new-learning on reconsolidation. Hippocampal and amygdala are differentially activated at the Final-test following New-learning administered during reconsolidation on Day 2, as given by the interaction term: (R_Day2_, _Exp_-R_Day3_, _Exp_) > (R_Day2_, _Ctrl_-R_Day3_, _Ctrl_); *P*-svc <0.05.

In a separate model (Intrusion model), the contrast “Intrusive Non-intrusive errors” revealed activation of the left inferior frontal gyrus (IFG, Fig. 4). This indicated the post-reactivation new-learning was associated with activation in the inferior frontal area, which has long been implicated in resolving interference between competing mnemonic representations of the originally learned and newly acquired associations (Badre, Poldrack et al. 2005). For consistency, we also performed SVC for the IFG using a functional mask defined in a previous study (mid-ventrolateral PFC, post-retrieval selection) (Badre, Poldrack et al. 2005) and confirmed the results (peak *P*-svc = 0.010).

**Figure 4.**
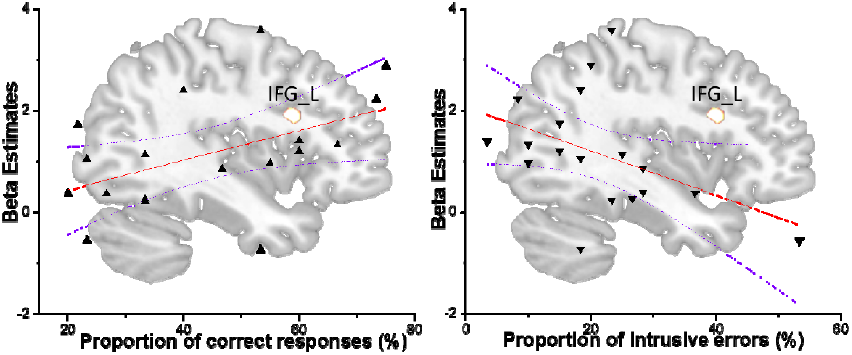
Engagement of IFG by new-learning intrusion. Left IFG activation measured on the Final-test reflects individuals’ variability in guarding against memory intrusion imposed by New-learning during reconsolidation on Day 2. The left inferior frontal gyrus is more activated by intrusive errors than by non-intrusive errors during Final-test (*P* <0.05). This difference in neural activation mediated the behavioral performance. Activation in the left IFG across participants is correlated positively with the percentage correct (*P* = 0.045, left), but is correlated in a negative trend with the number of intrusive errors (*P* = 0.060, right). Such IFG activation is however not correlated with the number of non-intrusive errors (*P* > 0.5, not shown). This result shows that post-reactivation new-learning manipulates memory by affecting reconsolidation on day 2, with the intrusion-effects being observed on day 3 in the Final-test. Triangles on the scatterplots represent individual subjects. The central line is the best linear fit with 90% confidence interval.

To elucidate the functional significance of the inferior frontal activation in relation to the behavioral results, we extracted the beta estimates of BOLD signal from the left IFG cluster and correlated these individual beta estimates with subjects’ percentage correct and intrusive proportion respectively. The individual beta estimates showed a significant positive correlation with the percentage correct rates (r = 0.48, *P* = 0.045, Fig. 4 left panel), whereas the beta estimates showed a negative correlation with the subjects’ intrusive proportion (r = −0.45, *P* = 0.060, Fig. 4 right panel). We interpret these pattern of results as that the IFG is involved in mediating the recollection bias towards the originally learned information. The more strongly the inferior frontal cortex is activated, the successful the participant would be in discriminating the respective memory traces associated with the original acquisition and new-learning, whereas a weaker inferior frontal involvement signifying a lower ability in dealing with the competition between mnemonic representations of the initially-learned and newly-acquired associations.

## 4 Discussion

In light of the previous studies which failed to observe the reconsolidation process in humans and non-human animals (Cammarota, Bevilaqua et al. 2004, Debiec, Doyère et al. 2006, Forcato, Argibay et al. 2009), we deduced several factors which might be instrumental for the reconsolidation processes at play. In declarative memories, content similarity shared between the acquisition and new-learning material is a key factor for effective intervention as only similar new materials were found to induce memory update, disruption or enhancement via reconsolidation (Forcato, Burgos et al. 2007, Hupbach, Gomez et al. 2007, Coccoz, Maldonado et al. 2011, Forcato, Rodriguez et al. 2011). Based on the results of the control experiment, we ascertain that the new-learning was most effective in affecting reconsolidation when “same faces” were employed. We thus assert that reconsolidation could be disrupted by post-reactivation new-learning *if and only if* the new material was similar enough to those involved in the acquisition, establishing content similarity in the associative memory traces between acquisition and new-learning to be a determinant factor. If the new-face-learning was distinct from the reactivated memory traces then these new-face-learning might have induced a different set of consolidation processes independently of the targeted reactivation.

In the rodents, any intervention disrupting memory reconsolidation is only effective when it is administered shortly after reactivation (Nader, Schafe et al. 2000, Debiec, LeDoux et al. 2002, Pedreira, Perez-Cuesta et al. 2002, Debiec and Ledoux 2004), suggesting that reconsolidation is a highly time-dependent phenomenon. In the humans, there has not been a consensus on the precise interval for this mnemonic fragility (Forcato, Burgos et al. 2007, Schiller, Monfils et al. 2010, Agren, Engman et al. 2012). Our current study incorporated a range of gradient-like post-reactivation delays. The New-learning administered within 20 minutes caused retrograde amnesia, whereas delays longer than that elicited no effect. Our results thus provide a qualifier on defining the critical time-window for post-reactivation manipulation to be effective for inducing forgetting: immediately after reactivation when memory is being updated. When the interval was long and beyond the susceptible period, the reactivated memories would become stable again and immune to any new-learning, thus no effect would be observed. This conclusion is further verified by the analyses of the intrusive effect reported in Fig. 2C, which illustrate that the differential intrusive effects induced by new-learning following different post-reactivation delays.

Our fMRI findings demonstrate how the memory systems might have acted interactively in declarative memory reconsolidation. It is known that memory reactivation will render consolidated memory (hippocampus-independent) to be hippocampus-dependent again (Debiec, LeDoux et al. 2002, Kelly, Laroche et al. 2003, Lee, Everitt et al. 2004). Our fMRI results reveal that memory processes during reconsolidation are hippocampus-dependent, strengthening the view that the hippocampal and amygdala involvement change with the passage of time during reconsolidation (Agren, Engman et al. 2012, Schwabe, Nader et al. 2012). When the post-reactivation manipulation requiring the hippocampus (and amygdala) to process new but similar information during active reconsolidation, the originally acquired memories would be affected by disruption or intrusion.

In contrast to previous studies (Nader, Schafe et al. 2000, Debiec and Ledoux 2004, Lee, Di Ciano et al. 2005), the amygdala activation was presently observed in the absence of emotional input or incentive factors (neutral faces location association). We proposed two possible explanations for this: First, the faces encoded by the participants might inherently carry emotional valence and collaterally engaged the amygdala. However, an alternative, more nascent, account is that the amygdala has a seat during declarative memories reconsolidation, irrespective of emotion aspects, acting in concert with the hippocampus. We are in favor of the latter account especially our results align with some recent causal evidence that the human amygdala possesses a general capacity to endogenously initiate memory prioritization processes of declarative memories without eliciting any subjective emotional response (Inman, Manns et al. 2018), establishing the amygdala as an overarching operator of downstream memory processes.

The activation in the left inferior frontal gyrus was differentially increased by intrusive events, suggesting that left IFG is involved in discriminating the originally learned and newly-learned memories and deciding which memories should be reactivated according to the cue (Zhang, Feng et al. 2004, Badre, Poldrack et al. 2005, Moss, Abdallah et al. 2005, St Jacques, Olm et al. 2013). Due to the high similarity between the originally learned and newly learned memories, the participants have to recollect the episodes in greater detail to overcome the competition and meet the goal in recalling the relevant, correct memories among competitive sources. In line with the view that the left ventral PFC mediates post-retrieval selection during source recollection and decision (Badre, Poldrack et al. 2005, Badre and Wagner 2007), our findings of increased left IFG activation characterize this region as a target area for manipulating memory retrieval especially during reconsolidation. The individual difference in left IFG activation among participants further serves as an indicator of individual’s ability in reconciling the mnemonic intrusion during memory reconsolidation.

## 5 Conclusion

Overall, we reveal three neuro-behavioral features in declarative memory reconsolidation in humans. The results provided insights into the mechanisms of episodic memory reconsolidation, suggesting that reactivation can indeed effectively trigger reconsolidation with several qualifiers. First, new-learning is effective only when sharing common components with initial learning (acquisition). Second, we establish the existence of a critical time-window for reconsolidation, defining it to be 20 minutes. Third, we show the involvement of the hippocampus and amygdala in integrating newly-formed memories during reconsolidation, and with the IFG resolving the mnemonic competition caused by the intrusion by newly-formed memories. From a translational perspective, the present findings support the possibility that non-invasive manipulation may one day make drug therapy obsolete and carry important implications for educational and clinical practices in devising learning strategies.

## Supplementary information

containing 2 figures and 3 tables is included.

## Author Contributions

All authors contributed to the study design. F. S., J. W., and Y. C. conducted the behavioral experiment. F. S. and J. L. performed the fMRI experiment. F. S., Y. K., Z. W., H. W. and S. C. K. analyzed the data. F. S., Z. W., H. W. and S. C. K. wrote and approved the final version of manuscript.

## Acknowledgements

We thank Qing Cai for her advice on MRI data collection and analysis.

## Open Practices Statement

The data that support the findings of this study are available from the corresponding author on request.

